# Acute and repeated intranasal oxytocin differentially modulate brain-wide functional connectivity

**DOI:** 10.1101/798025

**Authors:** Marco Pagani, Alessia De Felice, Caterina Montani, Alberto Galbusera, Francesco Papaleo, Alessandro Gozzi

**Author notes:** Shared co-first authorship (Marco Pagani, Alessia De Felice). Corresponding author: Alessandro Gozzi, Functional Neuroimaging Laboratory, Center for Neuroscience and Cognitive Systems @UniTn, Istituto Italiano di Tecnologia, Corso Bettini 31, 38068 Rovereto (TN), Italy.

## Abstract

Central release of the neuropeptide oxytocin (OXT) modulates neural substrates involved in socio-affective behavior. This property has prompted research into the use of intranasal OXT administration as an adjunctive therapy for brain conditions characterized by social impairment, such as autism spectrum disorders (ASD). However, the neural circuitry and brain-wide functional networks recruited by intranasal OXT administration remain elusive. Moreover, little is known of the neuroadaptive cascade triggered by long-term administration of this peptide at the network level. To address these questions, we applied fMRI-based circuit mapping in adult mice upon acute and repeated (seven-day) intranasal dosing of OXT. We report that acute and chronic OXT administration elicit comparable fMRI activity as assessed with cerebral blood volume mapping, but entail largely different patterns of brain-wide functional connectivity. Specifically, acute OXT administration focally boosted connectivity within key limbic components of the rodent social brain, whereas repeated dosing led to a prominent and widespread increase in functional connectivity, involving a strong coupling between the amygdala and extended cortical territories. Importantly, this connectional reconfiguration was accompanied by a paradoxical reduction in social interaction and communication in wild-type mice. Our results identify the network substrates engaged by exogenous OXT administration, and show that repeated OXT dosing leads to a substantial reconfiguration of brain-wide connectivity, entailing an aberrant functional coupling between cortico-limbic structures involved in socio-communicative and affective functions. Such divergent patterns of network connectivity might contribute to discrepant clinical findings involving acute or long-term OXT dosing in clinical populations.

## Introduction

Oxytocin (OXT) is a neuropeptide synthesized by magnocellular and parvocellular neurons of the paraventricular, supraoptic and accessory nuclei of the hypothalamus (Grinevich et al., 2016). OXT-producing neurons send widespread projections to brain areas involved in emotional, affiliative and socio-communicative functions in mammals (Ross and Young, 2009; Ferretti et al., 2019). Accordingly, central release of OXT results in a significant modulation of social recognition memory, social interactions, pair-bonding, emotion discrimination and maternal behavior across species (Neumann and Landgraf, 2012; Grinevich et al., 2016; Oettl et al., 2016; Johnson and Young, 2017; Ferretti et al., 2019). These properties have prompted research into the use of OXT as an adjunct treatment for developmental and psychiatric conditions characterized by social interaction deficits, such as autism spectrum disorders (ASD) (Harony and Wagner, 2010; Neumann and Landgraf, 2012; Lukas and Neumann, 2013; Tseng et al., 2014). Recent genetic and post-mortem studies have bolstered a putative etiological contribution of OXT deficits to ASD, fueling further interest in the therapeutic potential of OXT supplementation. For example, polymorphisms in the human OXT receptor (OXTR) gene are significantly associated with an ASD diagnosis and/or predict the severity of ASD (Tost et al., 2010; Parker et al., 2014). Similarly, post mortem difference in OXTR density in the brain of ASD and control patients have been recently reported, suggesting a potential involvement of the OXT system in complex social cognition in humans, including populations with ASD (Freeman et al., 2018).

Human investigations of the substrates and effects of OXT have been typically carried out via intranasal administration of the neuropeptide. This delivery route produces biologically effective concentrations in the central nervous system, minimizing systemic side effects and overcoming permeability limitations posed by the blood–brain barrier (Born et al., 2002; Lee et al., 2018). Several proof-of-concept investigations in humans have shown that single-dose (i.e. acute) intranasal administration of OXT in healthy adults increases trust, cooperation, and positive social interaction with respect to placebo (Kosfeld et al., 2005; Ditzen et al., 2009; Declerck et al., 2010; van Ijzendoorn et al., 2012). In keeping with these findings, task-based imaging studies have revealed that intranasal administration of OXT can functionally modulate brain regions involved in human socio-communication, including the amygdala, hippocampus, parahippocampal gyrus, medial prefrontal cortex, insula and caudate putamen (Striepens et al., 2012; Wittfoth-Schardt et al., 2012; Sripada et al., 2013; Hu et al., 2015; Kanat et al., 2015; Eckstein et al., 2017; Patin et al., 2018). Such an increased knowledge of the substrates modulated by OXT in the healthy brain have fueled clinical investigations of the therapeutic potential of this peptide in ASD. However, the studies published so far have produced conflicting results. Indeed, while multiple investigations have shown that *acute* intranasal OXT can improve some aspects of social functioning in ASD (Domes et al., 2007; Andari et al., 2010; Guastella et al., 2010; Auyeung et al., 2015), more extensive clinical testing via *repeated* daily dosing of OXT has so far failed to demonstrate substantial therapeutic or behavioral benefits (Anagnostou et al., 2012; Tachibana et al., 2013; Dadds et al., 2014; Guastella et al., 2015). While etiological heterogeneity of ASD and other experimental factors are likely to play a role in such clinical setbacks, the discrepant results obtained with acute or repeated OXT dosing raise the possibility that neuroadaptive mechanisms could bias the functional effects and neural response to exogenous OXT administration, a hypothesis partly supported by preliminary animal investigations (Bales et al., 2013; Rault et al., 2013; Huang et al., 2014).

Here we use functional magnetic resonance imaging (fMRI) to spatially resolve the neural circuitry and network systems engaged by acute and chronic (seven-day) dosing of OXT in the adult mouse brain. By leveraging covariance-based fMRI connectivity mapping (Schwarz et al., 2007a; Gozzi et al., 2010; Gozzi et al., 2012; Galbusera et al., 2017), we document that acute and repeated OXT administration elicits comparable evoked fMRI responses, but engage different patterns of cortico-limbic connectivity. Importantly, we also show that such connectional reconfiguration is accompanied by a paradoxical reduction in social interaction and communication in healthy wild type mice. Our results identify the neural circuitry engaged by exogenous OXT administration, and reveal a previously unreported homeostatic reconfiguration of functional connectivity upon extended OXT dosing in the mammalian brain.

## Materials and Methods

### Ethical statement

All *in vivo* experiments were conducted in accordance with the Italian law (DL 26/214, EU 63/2010, Ministero della Sanità, Roma). Animal research protocols were reviewed and consented to by the animal care committee of the Istituto Italiano di Tecnologia and the Italian Ministry of Health (authorization 560/2016PR to A. Gozzi). All surgical procedures were performed under anesthesia.

### Experimental animals

Male C57BL/6J mice (age range: 14-20 weeks) were housed, three to four per cage, in a climate-controlled animal facility (22 ± 2°C) and maintained on a 12-hour light/dark cycle, with food and water available *ad libitum*.

### OXT treatment

Mice were handled daily one week before the start of intranasal treatments with oxytocin or vehicle to reduce stress associated with the procedure. Oxytocin (OXT, Sigma Aldrich) was dissolved in distilled water and administered intranasally in a volume of 5 µl to each nostril, to deliver a total dose of 0.5 µg of peptide per mouse, corresponding to 0.3 International Units (UI). This dose is behaviorally active (Huang et al., 2014), elicits detectable fMRI responses in the mouse (Galbusera et al., 2017) and is allometrically comparable to OXT dosing used in human clinical studies, as discussed in great detail in (Galbusera et al., 2017). A schematic illustration of the employed treatment scheme is reported in Figure 1. Chronic intranasal treatment was carried out by administering the same dose of OXT twice a day for seven consecutive days, 12 hours apart (first treatment between 8-9 a.m., the second between 8 and 9 p.m.). Vehicle-treated control mice received the same volume of distilled water, acutely or chronically, for 7 days. Treatment groups consisted of 19 mice chronically treated with OXT (“OXT Chr”) and 17 control animals treated with vehicle. During imaging acquisition, each of these mice received a single challenge dose of OXT (2 µl per nostril to achieve a total dose of 0.5 µg per mouse) to elicit a detectable fMRI response that could be regionally quantified and be used in rCBV covariance mapping (Schwarz et al., 2007a). In the case of vehicle pre-treated animals, the OXT administered in the MRI scanner served as first acute OXT treatment, in an otherwise exogenous OXT naïve brain. For this reason we have termed this group “OXT”, referring to the fact that this animals only received a single dose of OXT (i.e. during the fMRI experiment). A separate group of 16 animals were chronically treated intranasally for seven consecutive days with vehicle (water, twice a day, 12 hours apart), and challenged intranasally inside the scanner with 2 µl of water per nostril (“OXT Chr”). This cohort (which we refer to as “Vehicle”) was employed as reference baseline control group with respect to which we assessed the effect of acute OXT treatment.

**Figure 1.**
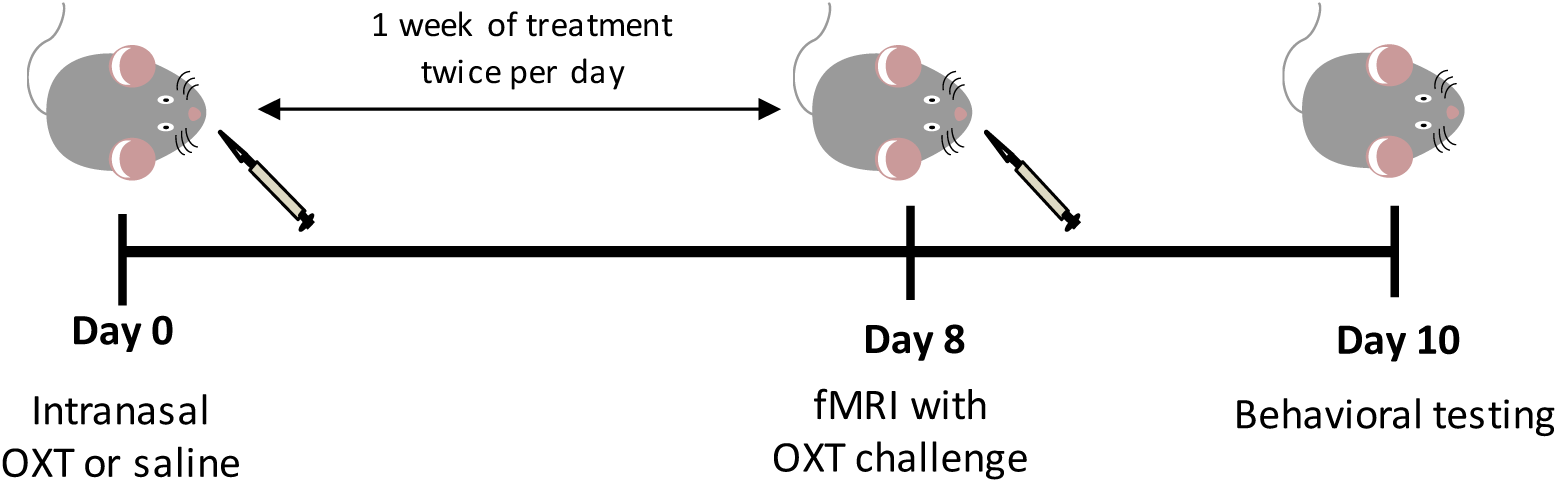
Timeline for fMRI and behavioral experiments. Mice were chronically treated with intranasal OXT or vehicle twice per day, 12 hours apart, for seven consecutive days (day 1-7). On day 8, mice underwent CBV-weighted fMRI experiments during which they received an intranasal OXT challenge. This challenge served as first OXT administration for the acute group. Three days later all mice were tested in a male-female social interaction test.

### Functional Magnetic Resonance Imaging (fMRI)

Animal preparation for fMRI has been described in great detail elsewhere (Sforazzini et al., 2014; Pagani et al., 2019). The protocol utilized is optimized for physiological stability and permits monitoring of peripheral blood pressure and arterial blood gases (Ferrari et al., 2010; Liska et al., 2015; Bertero et al., 2018). Briefly, the left femoral artery was cannulated under deep isoflurane anesthesia, and mice were imaged under halothane sedation (0.9%). Mechanical ventilation following endotracheal intubation was used to avoid hypercapnia and maintain arterial blood gas values within physiological range (pCO_2_ < 40 mmHg, pO_2_ > 90 mmHg). Functional MRI time series were acquired with a 7T Pharmascan (Bruker, Biospin), using a 72 mm birdcage transmit coil and a 4-channel solenoid coil for signal reception. For each session, *in vivo* anatomical images were acquired with a fast spin echo sequence (TR = 5500 ms, TE = 60 ms, matrix size 192 × 192, FOV 2 × 2 cm, 24 coronal slices, slice thickness 500 µm). Co-centered fMRI time series were acquired using a FLASH sequence with the following parameters: TR = 288 ms, TE = 3.1 ms, a = 30°; 180 × 180 × 600 µm resolution, dt = 60 s, NR = 50 corresponding to 50 min total acquisition time. Twenty minutes before intranasal administration of OXT, fMRI images were sensitized to reflect alterations in relative cerebral blood volume (rCBV) using 5 µl/g of blood-pool contrast agent (Molday Ion, Biopal, Worcester, MA, USA) as previously described (Galbusera et al., 2017).

To allow for the intranasal administration of OXT or vehicle inside the scanner, an intranasal catheter was inserted into each nostril as previously described. Briefly, an Hamilton Syringe 710N 100 µl was coupled with two PTFE cannulas 1 Fr (one for each nostril) using a 60 cm-long PE10 catheter connected to a sterile Y connector (22ga). Standard silicon laboratory tubing was used for all coupling connections. The syringe was then filled with distilled water. Four microliters of air (2 µl per each end of the intranasal cannula) were sucked into the cannulas to avoid mixing with the peptide solution to be administered. To avoid incidental pre-administration solution leakage, two additional microliters of air were also sucked in the terminal end of the cannula after compound loading. The two prefilled cannulas were then inserted (9 mm) into the nostrils.

### rCBV quantification and functional connectivity analysis

Regional fMRI response to the OXT challenge was quantified in the form of relative CBV (rCBV) in regions of interest as previously described (Galbusera et al., 2017). Prior rCBV quantification, fMRI time series were motion corrected and spatially normalized to a common reference space. A neuroanatomical parcellated atlas was then co-registered to the template (Pagani et al., 2016b) and used as reference to locate volumes of interest (VOIs) to quantify regional rCBV response. OXT-induced changes in connectivity were mapped using group-level rCBV-covariance mapping owing to the exquisite sensitivity of this approach to neuromodulatory induced changes in functional connectivity (Schwarz et al., 2007b; Gozzi et al., 2010), and its superior ability to resolve with high spatial resolution neurotransmitter systems (Schwarz et al., 2007b; Gozzi et al., 2010).

To assess whether acute and chronic OXT treatment results in differential re-organization of functional networks, we carried out seed based analysis to map target regions of long-range functional connectivity. This method relies on the computations of dependencies of rCBV changes between brain regions (i.e. covariance mapping) in response to a pharmacological challenge as previously described (Gozzi et al., 2010; Gozzi et al., 2012; Razoux et al., 2013). Specifically, the rCBV response in anatomical volumes of interest (VOIs) was used as a regressor to generate voxel-wise correlational maps and identify voxels that positively correlate with the seeding VOIs (Pagani et al., 2016a). For each treatment cohort, the resulting functional connectivity maps were then thresholded at *t* > 2.7 (*p* < 0.01) and familywise error (FWE) corrected for multiple comparisons with a cluster defining threshold of *p* = 0.05 (Worsley et al., 1992) as implemented in FSL (Jenkinson et al., 2012). Voxel-wise treatment-dependent intergroup differences (i.e. OXT vs. vehicle and OXT chr vs. OXT) were then assessed using a permutation-based unpaired Student’s t test, thresholded at *p* > 0.05 and FWE corrected for multiple comparisons with a cluster defining threshold of *p* = 0.05. To quantify intergroup differences in rCBV functional connectivity, we also carried out a treatment-dependent slope analysis by calculating rCBV Pearson’s correlation between the seeds and representative target regions (GraphPad v8.0).

### Behavioral tests

Three days after the imaging studies, animal belonging to the “OXT Chr” and “OXT” group mice underwent a male-female social interaction testing as previously described (Scattoni et al., 2011; Huang et al., 2014; Michetti et al., 2017; Liska et al., 2018). Briefly, socio-communicative testing was conducted in lightly illuminated (5 ± 1 lux) 2150E Tecniplast cages (35.5 × 23.5 × 19cm). A Canon LEGRIA HF R806 digital camera was used to record the testing session. Before behavioral testing, each male mouse was placed in the test cage and left to habituate for one hour. Then, an unfamiliar female in estrus was placed into the testing cage for a 5-min test session. The vaginal estrus condition of each female was assessed as previously described (Rugh, 1990). Scoring of social investigation was conducted by an observer blind to mouse treatment, and multiple behavioral responses exhibited by the test mouse were measured, including anogenital sniffing (direct contact with the anogenital area), body sniffing (sniffing with the flank area), head sniffing (sniffing with the head area) and following (time spent in following the female mice). Social investigation was defined as the sum of duration of total sniffing and following behaviors and expressed in seconds. During male-female social interaction test, we also measured ultrasonic vocalizations (USVs) emitted by male mice with an Avisoft UltraSoundGate condenser microphone capsule CM16 (Avisoft Bioacoustics, Berlin, Germany) mounted 20 cm above the cage and we recorded USVs using RECORDER (v3.2). Settings included sampling rate at 250 kHz; format 16 bit. The ultrasonic microphone was sensitive to frequencies between 10 and 180 kHz. For acoustical analysis, recordings were exported to Avisoft SASLab Pro (v4.4) and a fast Fourier transform was conducted as previously described (Scattoni et al., 2011). USVs were quantified as number of recorded events and expressed as frequency.

## Results

### Comparable fMRI reactivity in mice acutely or chronically exposed to intranasal OXT

Previous research has hinted at a possible decreased neural responsivity to repeated administration of OXT (Huang et al., 2014). To probe whether chronic OXT administration results in decreased functional activity *in vivo*, we quantified and compared the rCBV response produced by acute (OXT group) and chronic (OXT Chr group) intranasal OXT administration. In keeping with previous fMRI investigations (Galbusera et al., 2017), temporal profiles of rCBV revealed the presence of sustained rCBV responses in hippocampal (Figure 2A, top, t_29_ = 23.2, *p* < 0.001) and forebrain areas rich in OXT receptors, such as the diagonal band (Figure 2A, bottom, t_29_ = 17.9, *p* < 0.001). Interestingly, fMRI responsivity to OXT challenge appeared to be comparable in animals acutely or chronically exposed to the peptide (Figure 2B, top, t_29_ = 18.2, *p* < 0.001, bottom, t_29_ = 17.5, *p* < 0.001). This observation was corroborated by a formal quantification of OXT-evoked fMRI responses across a wider set of cortical and subcortical regions, in which comparably large fMRI responses were observed across groups when quantified either at the voxel-level, or in volumes of interests (Figure 2C, *p* > 0.31, all regions).

**Figure 2.**
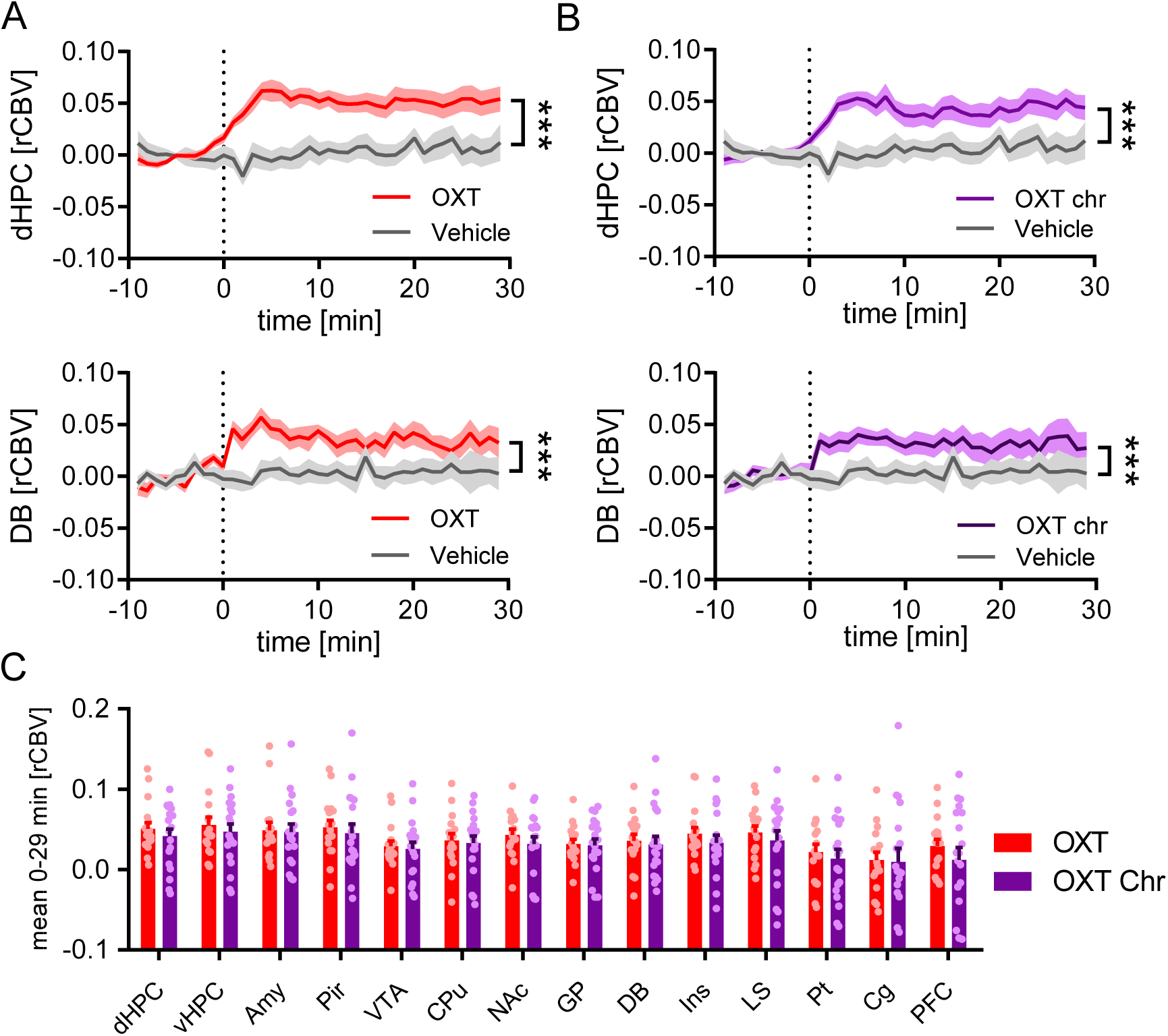
Comparable fMRI reactivity in acutely and chronically OXT-treated mice. OXT-elicited fMRI timecourses in the dorsal hippocampus and diagonal band of mice acutely (A, OXT, red) and chronically (B, OXT Chr, purple) exposed to OXT. Mice were challenged with OXT or vehicle at time 0. (C) rCBV quantification in neuroanatomical volumes revealed comparable fMRI reactivity in OXT chr and OXT mice. dHPC, dorsal hippocampus; vHPC, ventral hippocampus; Amy, amygdala; Pir, piriform cortex; VTA, ventral tegmental area; CPu, caudate putamen; NAc, nucleus accumbens; GP, globus pallidus; DB, diagonal band; Ins, insula; LS, lateral septum; Pt, parieto-temporal cortex; Cg, cingulate cortex; PFC, prefrontal cortex. Data are plotted as mean ± SEM for each experimental group. ***p < 0.001.

Importantly, arterial blood pressure recordings showed that mean blood pressure was within the physiological range of auto-regulation (80-120 mmHg) throughout the recording sessions in both groups of subjects (Gozzi et al., 2007), ruling out putative peripheral vascular contributions to the central hemodynamic effects observed. Together, these results suggest that acute and chronic OXT treatment elicit comparable regional functional activation (e.g. *fMRI reactivity*) in the mouse brain.

### Acute intranasal OXT increases functional connectivity in cortico-limbic region

To map the network substrates modulated by acute OXT in the mouse brain, we carried out covariance-based inter-regional rCBV mapping in representative cortical and subcortical areas using voxel-wise seed based correlations (Schwarz et al., 2007b). Because OXT is synthesized in the hypothalamus and is released to distal cortico-limbic regions through widespread neuronal projections (Ross and Young, 2009; Grinevich et al., 2016; Ferretti et al., 2019), we first probed the functional connectivity between this region and its long-range targets in mice acutely challenged with OXT compared to vehicle treated controls. This analysis revealed a significant increase in functional connectivity between the hypothalamus and the amygdala, ventral hippocampus and nucleus accumbens (Figure 3A, *t* test, *p* < 0.05, FWER cluster-corrected at *p* < 0.05). To further dissect the functional circuitry engaged by OXT, we next probed some of these over-connected targets using an additional set of seed areas (Figure 3B-C). Functional connectivity mapping of the amygdala revealed increased coupling with the ventral tegmental area and postero-ventral hippocampal areas (Figure 3B, *t* test, *p* < 0.05, FWER cluster-corrected with *p* < 0.05), whilst functional probing of the ventral hippocampus showed hyper-connectivity with thalamic regions (Figure 3C, *t* test, *p* < 0.05, FWER cluster-corrected with *p* < 0.05). Further seed-based mapping of the prefrontal cortex, revealed foci of significant long-range hyper-connectivity between this region and the periaqueductal grey (Figure 3D, *t* test, *p* < 0.05, FWER cluster-corrected, with cluster defining threshold of *p* < 0.05). Collectively, these results document that acute OXT administration enhances functional connectivity between limbic and prefrontal regions areas that are key for social functioning and behavior.

**Figure 3.**
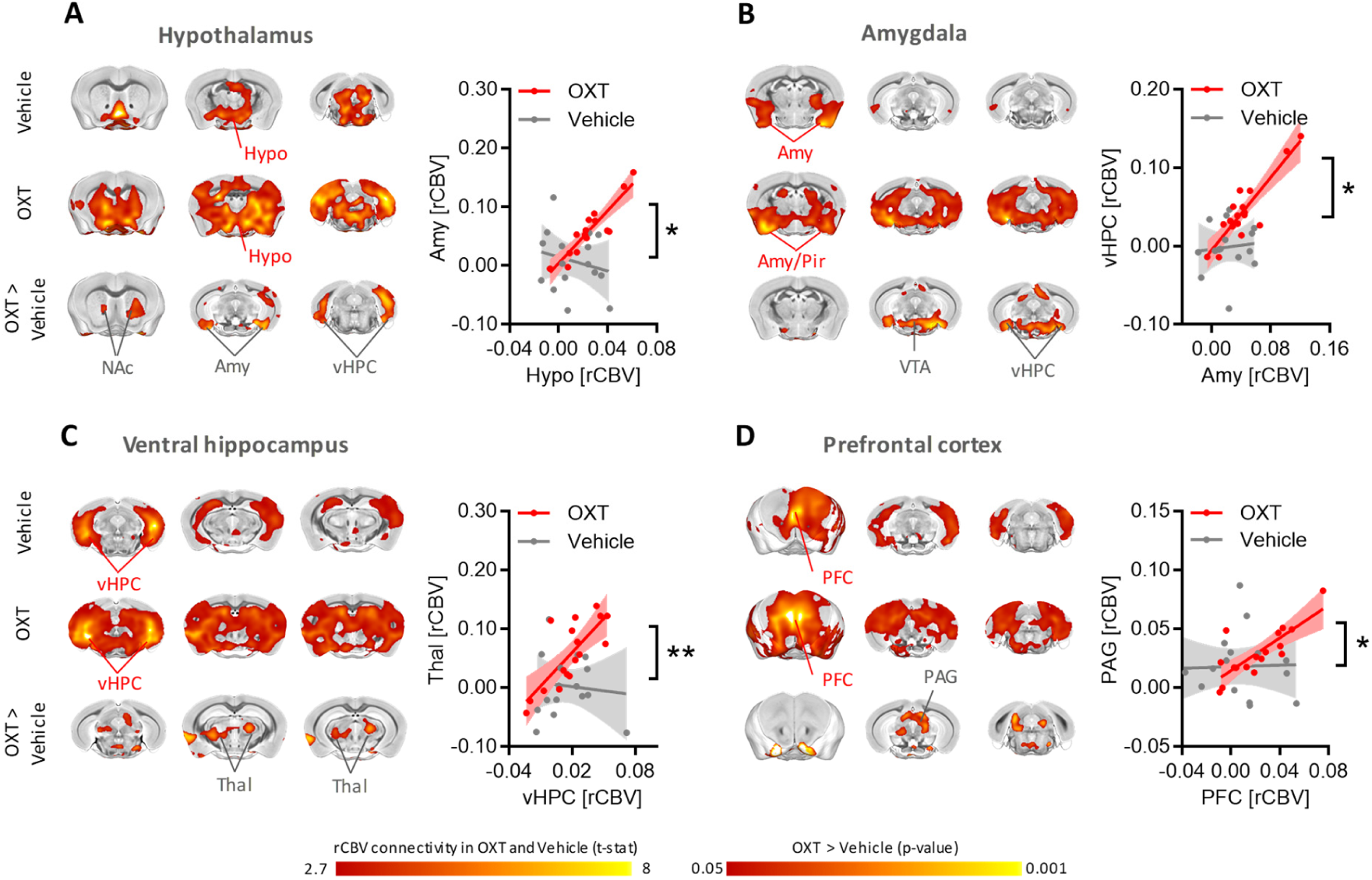
Acute OXT administration increases connectivity between cortico-limbic areas. (A) Spatial extension of brain regions exhibiting significant functional connectivity with the hypothalamus (seed, red lettering) in mice challenged with vehicle (vehicle, top row) or acute OXT (OXT, middle row). Red-yellow represents brain areas of significant correlation (i.e. functional connectivity) with the hypothalamic seed. Intergroup treatment-dependent analysis (OXT > vehicle, bottom row) revealed significantly increased functional connectivity between hypothalamus and ventral hippocampus, amygdala and nucleus accumbens in OXT-treated mice as compared to vehicles. Here, red-yellow represent brain regions exhibiting significantly increased connectivity with the hypothalamus in OXT-treated vs. vehicle mice. Slope analysis (right panel) confirmed increased functional connectivity between hypothalamus and amygdala in OXT-vs. vehicle-treated mice. Seed based mapping also revealed increased functional connectivity between (B) amygdala and ventral hippocampus and VTA, (C) the ventral hippocampus and thalamus and (D) the prefrontal cortex and periaqueductal grey. Scatterplots quantifies functional connectivity (slope) between the seed and a representative brain region in OXT treated (OXT, red line) as compared to vehicle (vehicle, grey line) mice. Shaded areas represent SEM. Amy, amygdala; NAc, nucleus accumbens; Hypo, hypothalamus; PAG, periacqueductal gray; Pir, piriform cortex, PFC, prefrontal cortex; Thal, thalamus; vHPC, ventral hippocampus; VTA, ventral tegmental area. *p < 0.05, **p < 0.01.

### Repeated OXT treatment reconfigures brain-wide functional connectivity

To assess whether repeated OXT administration would produce neuroadaptive changes in functional connectivity, we next probed covariance mapping in mice chronically treated with OXT, and mapped global functional network changes with respect to mice receiving an acute OXT challenge. Notably, and in contrast with our quantifications of rCBV responsivity, seed based connectivity mapping in OXT Chr mice revealed qualitatively different patterns of connectivity, characterized by a largely more widespread spatial extension of the probed network, and loss of network specificity, suggestive of a possible connectional reconfiguration. These features were apparent when qualitatively assessed in three experimental groups of this study (vehicle, OXT, and OXT Chr, respectively) using circular plots depicting the strongest (Pearson’s r > 0.85) inter-regional functional connections (Figure 4).

**Figure 4.**
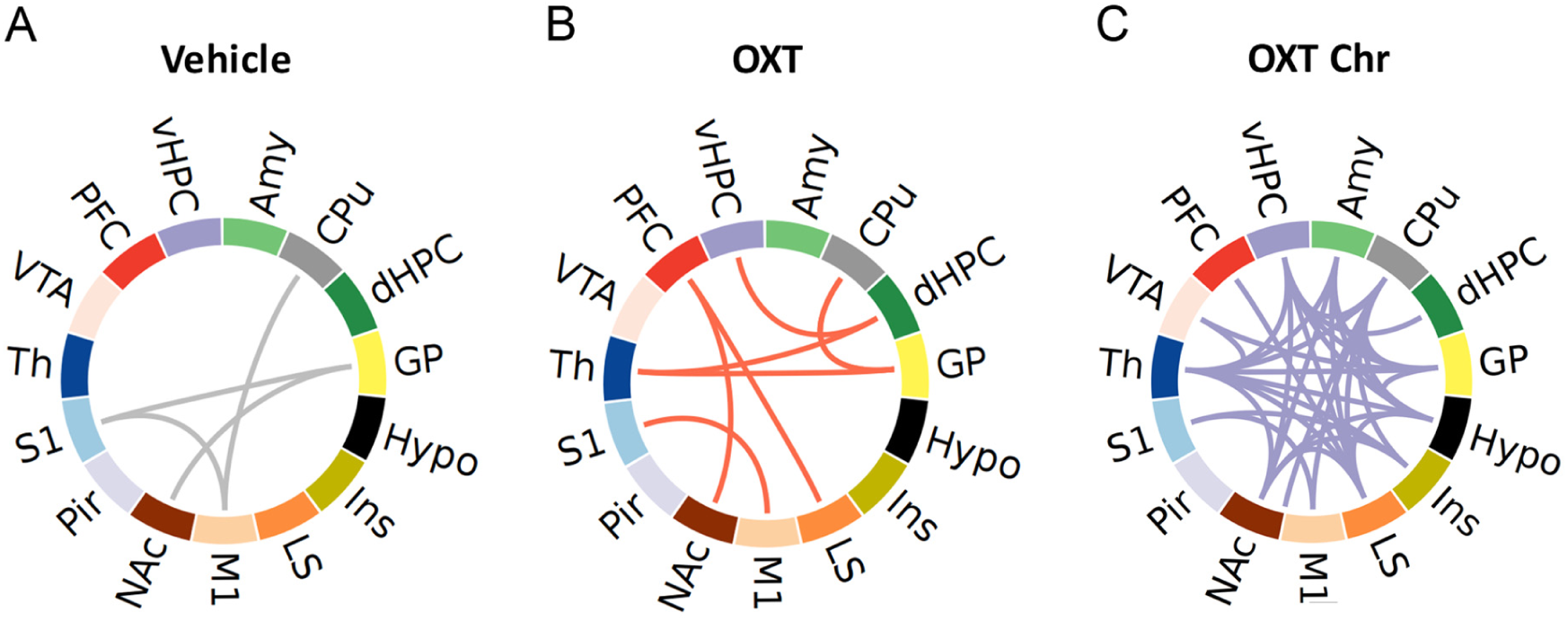
Circular plots of fMRI connectivity as a function of treatment. Circular layouts depicting interregional rCBV functional connectivity in mice treated with (A) vehicle, (B) acute OXT or (C) chronic OXT. Each link represents functional connectivity between regions exhibiting a Pearson’s correlation greater than 0.85. Amy, amygdala; Cpu, caudate putamen; dHPC, dorsal hippocampus; GP, globus pallidus; Hypo, hypothalamus; Ins, insula; LS, lateral septum; M1, primary motor cortex; NAc, nucleus accumbens; Pir, piriform cortex; S1, primary somatosensory cortex; Th, thalamus; VTA, ventral tegmental area; PFC, prefrontal cortex; vHPC, ventral hippocampus.

Voxel-wise seed based probing of functional connectivity differences between acute and chronic OXT dosing corroborated these findings, revealing bilateral foci of increased connectivity between the prefrontal cortex and amygdaloid/piriform areas, (Figure 5A, *t* test, *p* < 0.05, FWER cluster-corrected at *p* < 0.1), and between the amygdala and large somato-motor cortical territories (Figure 5B, *t* test, *p* < 0.05, FWER cluster-corrected at *p* < 0.1). In keeping with these findings, seed-based probing of somatosensory cortices revealed reciprocally increased functional connectivity with the amygdala and piriform areas (Figure 5C, *t* test, *p* < 0.05, FWER cluster-corrected, *p* < 0.1). Finally, we also found the ventral tegmental area be over-connected with the ventral hippocampus, habenula, and periaqueductal grey in OXT treated mice compared to control OXT-naïve subjects (Figure 5D, *t* test, *p* < 0.05, FWER cluster-corrected, *p* < 0.1), while no area of increased functional connectivity was instead observed between the hypothalamus and its regional targets (data not shown). These results show that repeated OXT administration reconfigures brain-wide functional coupling between cortico-limbic and somatosensory cortical regions, resulting in widespread patterns of overconnectivity.

**Figure 5.**
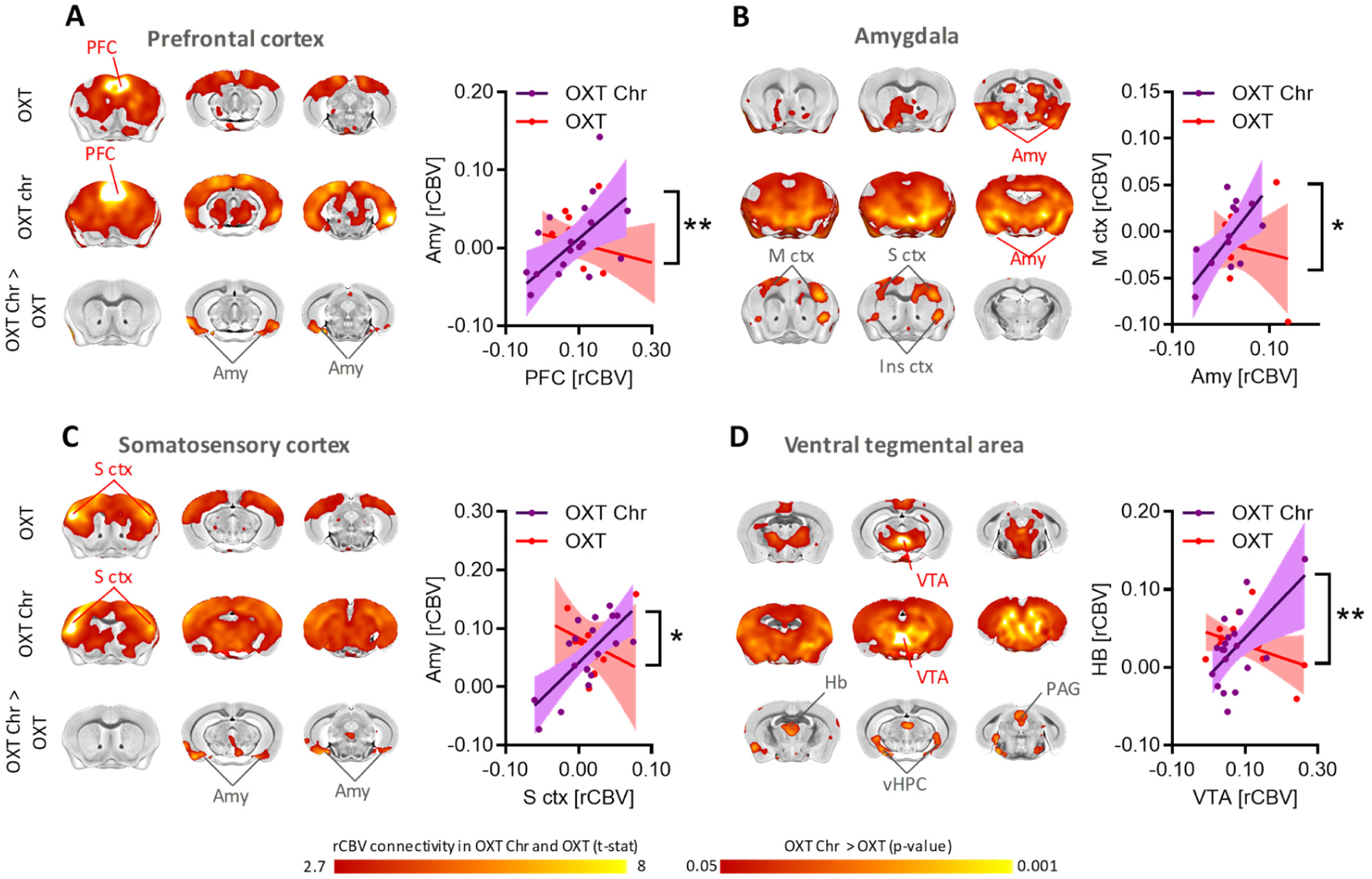
Chronic OXT treatment results in a reconfigured pattern of connectivity. (A) Spatial extension of brain regions exhibiting significant functional connectivity with the prefrontal cortex (seed, red lettering) in mice treated with acute (OXT, top row) or chronic OXT (OXT chr, middle row). Red-yellow represent brain areas of significant correlation (i.e. functional connectivity) with prefrontal seed region. Chronic OXT enhanced functional connectivity between prefrontal cortex and amygdala (A), somato-motor cortices and amygdala (B-C) and the VTA with habenula, ventral hippocampus and periaqueductal grey (D). Scatterplots quantify functional connectivity (slope) between the seed and a representative area for chronically (OXT chr, purple line) and acutely (OXT, red line) treated mice. Shaded areas represent 95% Confidence interval. Amy, amygdala; M ctx, motor cortex; S ctx, somatosensory cortex; LS, lateral septum; PAG, periaqueductal gray; PFC, prefrontal cortex, vHPC, ventral hippocampus. VTA, ventral tegmental area. *p < 0.05, **p < 0.01.

A schematic description of the results obtained with seed-based probing is reported in Figure 6, in which we depict the substrates engaged by acute OXT administration (with respect to baseline vehicle, Figure 6A) and by chronic OXT administration (with respect to acute OXT, Figure 6B), respectively.

**Figure 6.**
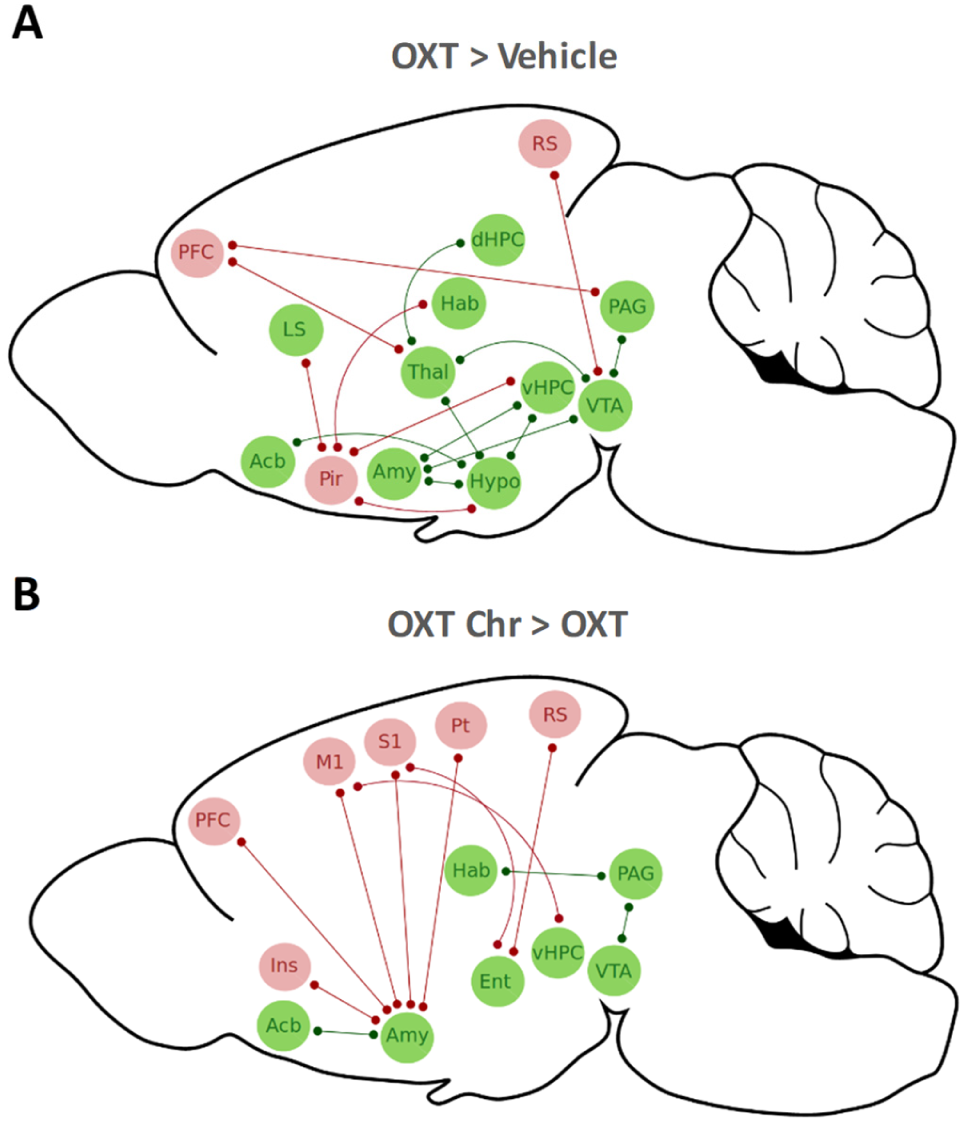
Brain-wide circuitry engaged by acute and chronic OXT administration. (A) Aggregated description of cortical (pink) and subcortical (green) substrates exhibiting increased functional connectivity in mice acutely treated with OXT with respect to baseline vehicle, as assessed with seed based mapping. Only links between areas showing significant intergroup functional connectivity changes are reported (p < 0.05). (B) Mice chronically treated with OXT exhibit a more widespread increase in functional connectivity, critically involving the amygdala and extended cortical brain regions. Acb, nucleus accumbens; Amy, amygdala; dHPC, dorsal hippocampus; Ent, entorhinal cortex; Hab, habenula; Hypo, hypothalamus; Ins, insular cortex; LS, lateral septum; M1, primary motor cortex; PFC, prefrontal cortex; Pir, piriform; Pt, parieto-temporal cortex; RS, retrosplenial; S1, primary somatosensory cortex; Thal, thalamus; PAG, periaqueductal gray; Pir, piriform cortex; vHPC, ventral hippocampus; VTA, ventral tegmental area.

### Chronic OXT dosing impairs socio-communicative behavior

Finally, to probe the behavioral relevance of the mapped connectional changes, we assessed the effect of chronic (7 week, daily administration) vs. acute OXT administration on social and communicative functions using a male-female interaction test in the same mice employed in the fMRI studies (Figure 7). Interestingly, treatment-dependent behavioral quantifications showed a prominent reduction of social behaviors (Figure 7A, t test, t = 2.17, p = 0.019), along with a trend for reduced USVs emission (Figure 7B, t test, t = 1.78, p = 0.087) in mice chronically treated with OXT compared to acute OXT controls, replicating previous observations of impaired social function in mice receiving extended OXT dosing (Huang et al., 2014). These findings suggest that aberrant functional over-connectivity associated with chronic OXT dosing might detrimentally affect higher order socio-communicative functions in healthy mice.

**Figure 7.**
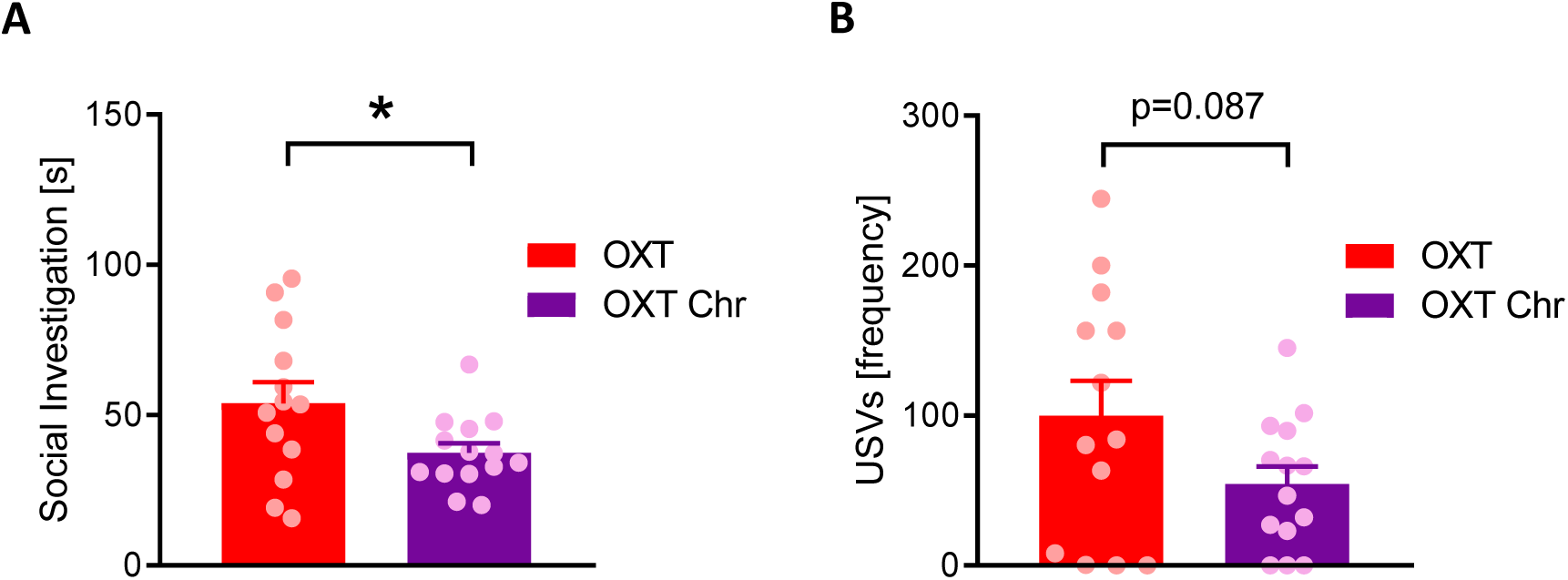
Chronic OXT treatment impairs social and communicative behaviors in mice. A) Male-female social interaction test revealed impaired social investigation in mice repeatedly treated with OXT with respect to mice that received acute OXT treatment. (B) Concomitant vocalization recordings revealed reduced frequency of USVs emitted by mice chronically vs. acutely treated with OXT. Bar graph represents mean ± SEM. ^*^p<0.05.

## Discussion

Intranasal administration of OXT activates neural substrates involved in socio-affective behavior and it is currently explored as an adjunctive treatment for ASD and related neurodevelopmental disorders characterized by social interaction deficits. Here we document that acute OXT dosing enhances connectivity in mouse brain regions involved in socio-communicative processing. We also show that repeated administration of this neuropeptide leads to a substantial reconfiguration of this pattern of connectivity, encompassing an exacerbation of cross-regional coupling and aberrant hyper-connectivity between amygdala and wide motor-sensory cortical areas.

Previous human studies have probed the substrates targeted by acute OXT administration via task-based protocols engaging brain areas involved in processing of social and cognitive stimuli, such as the amygdala (Kirsch et al., 2005; Domes et al., 2007; Gamer et al., 2010; Bethlehem et al., 2013). While these investigations have proven useful in elucidating some of the circuit elements modulated by this neuropeptide, task-based approaches are sensitive only to changes occurring in the brain areas engaged by the task employed. To overcome these limitations, attempts to map the regional targets of OXT via brain-wide task-free brain mapping have been described. Using pharmacological fMRI, (i.e. the use of fMRI to map drug-elicited hemodynamic responses; (Gozzi et al., 2008)), the ability of OXT to activate basal forebrain and ventral striatal areas involved in social behaviors has been demonstrated in human (Paloyelis et al., 2014) and mouse (Galbusera et al., 2017). These studies have recently been complemented by initial investigations of the effect of OXT on interregional synchronization, or “functional connectivity”, between brain areas. By using resting state fMRI, age-by-sex dependent strengthening of amygdala - prefrontal connectivity has been reported upon acute OXT administration (Sripada et al., 2012; Ebner et al., 2016). Other investigators have expanded these observations, reporting increased functional coupling between cortico-striatal regions (Bethlehem et al., 2017), and bidirectional changes in connectivity of the ventral attention networks with cingulate-opercular and default mode network (Brodmann *et al*, 2017). By contrast, chronic OXT administration has been reported to increase functional connectivity between the amygdala and dorsal anterior cingulate cortex in healthy volunteers (Kovács and Kéri, 2015), and between the anterior cingulate and medial prefrontal cortex in ASD patients (Watanabe *et al*, 2015). Despite these encouraging results, inconsistencies exist about the specific substrates and networks modulated by this neuropeptide (Anagnostou et al., 2014), and a number of investigators have failed to identify any significant OXT-induced modulation (Fan *et al*, 2014; Riem *et al*, 2013), questioning the robustness of some of these initial findings. Moreover, rigorous comparisons of the effects of acute and repeated OXT dosing on brain-wide function and connectivity are lacking.

Our study address these questions by using group-level rCBV covariance connectivity measurements in the mouse. Compared to standard rsfMRI mapping (Sforazzini et al., 2014; Gozzi and Schwarz, 2016; Bertero et al., 2018), covariance fMRI is exquisitely sensitive to neuromodulatory effects, and allows to resolve the connectivity patterns of subcortical nuclei with a spatial resolution that at present does not appear to be attainable with resting state fMRI (Schwarz et al., 2007b; Schwarz et al., 2007a; Schwarz et al., 2007c; Gozzi et al., 2010; Gozzi et al., 2012). Supporting the specificity of our findings, the observed pattern of increased functional connectivity upon acute OXT dosing encompasses nuclei and brain regions innervated by OXT hypothalamic neurons and pivotally involved in social memory and socio-communicative functions, such as the amygdala and olfactory cortices, nucleus accumbens, ventral hippocampus and prefrontal cortex (Goodson and Kingsbury, 2013). Interestingly, our mapping also highlighted enhanced coupling between the prefrontal cortex, the periaqueductal grey and the thalamus, a neural pathway implicated in socially directed activity (Stoop, 2014; Benekareddy et al., 2018), cognition (Bicks *et al*, 2015), as well as passive and active defensive responses to innate stimuli (Brandão et al., 2008; Nakajima et al., 2014; Ko, 2017), a set of behavioral functions modulated by endogenous OXT (Neumann and Landgraf, 2012). Taken together, these observations suggest that exogenous OXT can strengthen the functional coupling between the hypothalamus (where OXT-producing neurons reside) and several of its main target regions, hence reinforcing a distributed neural circuitry that is critical for social recognition and affective behavior (Neumann and Landgraf, 2012; Haak et al., 2018).

Interestingly, the brain-wide patterns of functional connectivity identified upon repeated OXT dosing were qualitatively different from those observed with acute treatment, hinting at the presence of neuroadaptive mechanisms leading to a reconfiguration of inter-regional coupling. A notable feature of this reconfigured pattern of connectivity was its widespread spatial distribution, indicative of reduced network specificity, together with an especially prominent involvement of the amygdala and basal olfactory cortices, which appeared to be strongly coupled to large motor sensory cortical regions. Notably, the same animals displayed impaired social behavior in a male-female social interaction, replicating previous observations of diminished social activity in chronically dosed rodents (Bales et al., 2013; Huang et al., 2014). The amygdala plays a key contribution in the processing of emotional and social relevant stimuli (Ferretti et al., 2019) and, in rodents, visual and auditory signals as well as input from the main and accessory olfactory bulbs converge directly to this region, where they are integrated and parsed based on their social relevance (Choleris *et al*, 2009). These functions suggest a view in which the social impairments observed in the chronic OXT group could therefore be the expression of an aberrant and/or overly distributed functional coupling between amygdaloid-olfactory regions and motor-sensory areas, resulting in a defective multi-sensory integration which could in turn bias the ensuing behavioral responses (Chen and Hong, 2018). In addition to this, we note that chronic OXT administration also exacerbated the functional coupling of the whole mesolimbic dopamine system, a pathway critical for the control of the reinforcing and aversive value of social stimuli (Dölen et al., 2013; Hung et al., 2017). It is therefore possible that imbalances in the tuning of social valence could also contribute to the observed behavioral impairment.

Additional factors, unrelated to or independent of inter-regional coupling, could also play a role, alone, or in combination, in the divergent behavioral profile observed between acute and repeated OXT dosing. One possibility is the presence of a “priming” effect, according to which endogenous OXT synthesis is overstimulated by the exogenous administration (Ludwig and Leng, 2006). Other authors have implicated neuroadaptive changes in NMDA-based glutamatergic transmission in the prefrontal cortex upon repeated OXT dosing in rodents (Benner et al., 2018), or called into play OXT receptor downregulation and desensitization (Huang et al., 2014), a hypothesis that however does not appear to be supported by our evidence of unaltered rCBV responsivity in acutely or repeatedly dosed animals. Further studies are required to characterize the precise mechanisms underlying the neuroadaptational changes we mapped here.

While caution must be exercised when extrapolating rodent findings to clinical populations, the remarkable functional reconfiguration we did observe warrants a discussion of the possible clinical implication of our result. It is indeed interesting to note that highly promising initial investigations of the acute effect of OXT in control populations (Meyer-Lindenberg et al., 2011) and patients with ASD (Andari et al., 2010; Zink and Meyer-Lindenberg, 2012; Gordon et al., 2013), have so far failed to translate into consistent, clinically significant endpoints when the therapeutic potential of OXT has been rigorously assessed in prolonged administration regimens (Leng and Ludwig, 2016; Yatawara et al., 2016; Yamasue et al., 2018). The heterogeneity of ASD and our inability to stratify patient population, together with the presence of robust placebo effects, are all likely primary contributors to such inconsistent clinical findings (Leng and Ludwig, 2016; Yamasue et al., 2018). Nevertheless, our observation that repeated OXT dosing could behaviorally and functionally bias the effects of this peptide should not be neglected, as similar hemostatic mechanisms could occur in human populations, and could represent a possible confounding factor when proof of concept investigations are to be translated into rigorous therapeutic testing. The design of ad hoc studies in which the acute and chronic effect of OXT are rigorously compared using socio-behavioral and imaging readouts in humans seems like an easily attainable goal for future clinical studies aimed at probe the actual translational relevance of our findings.

In conclusion, we describe the brain-wide circuitry engaged by acute and repeated OXT administration, and show that repeated dosing exacerbates cortico-limbic and dopamine mesolimbic connectivity, leading to impaired socio-behavioral functions. Our results shed light on the functional neurocircuitry engaged by OXT, and the neuroadaptive brain-wide response elicited by prolonged OXT administration.

## Acknowledgments

A.G. acknowledges funding by the Simons Foundation (SFARI 400101, A. Gozzi), the Brain and Behavior Foundation (2017 NARSAD, Independent Investigator Grant 25861), the NIH (1R21MH116473-01A1) and European Research Council (ERC, DISCONN, Grant Agreement 802371). M.P. is supported by European Union’s Horizon 2020 research and innovation programme (Marie Sklodowska-Curie Global Fellowship - CANSAS, GA845065).

